# Functional potential of bacterial strains in the premature infant gut microbiome is associated with gestational age

**DOI:** 10.1101/530139

**Authors:** Sumayah F. Rahman, Matthew R. Olm, Michael J. Morowitz, Jillian F. Banfield

## Abstract

The gut microbiota of premature and full-term infants have many known differences, but the extent to which the degree of prematurity influences the structure and functional potential of the microbiome has not been deeply explored. Here, we used genome-resolved metagenomics to address how gestational age impacts the premature infant gut microbiome. We found that gestational age is associated with species richness, with more premature infants having lower species richness; this effect lasts until the fourth week of life. Novel *Clostridium* species and strains related to *Streptococcus salivarius* and *Enterococcus faecalis* colonize infants of different gestational ages, and the metabolic potential of these organisms can be distinguished. Thus, we conclude that the extent of prematurity, or directly linked factors, can be an important influence on the microbiome and its functions.

## Introduction

The human gut microbiome plays many important roles, including the extraction of nutrients from food, metabolizing toxins, immunomodulation, and protection from pathogens^1^. Infants, near-sterile when born, obtain microbes from their mother and their environment^2–4^. The gut microbiome of infants is known for its simplicity and low complexity compared to the gut microbiome of older children and adults^5^. Premature infants, born before they have reached 37 weeks *in utero*, harbor gut microbial communities of even lower complexity than full-term infants, as they are colonized by tenfold fewer species^6^. Premature infant gut microbiomes display abrupt shifts in composition^7^, and may have a different taxonomic makeup than microbiomes of full-term infants^6,8,9^. Studies on premature infants suggest that the extent of the infant’s prematurity influences their gut microbiome. A recent study, utilizing 16S rRNA gene sequencing, revealed that bacterial alpha diversity varies based on the infants’ gestational age, with more premature infants having less diverse microbiomes. This study also showed that infants born at a later gestational age had greater abundance of *Bifidobacterium* and *Streptococcus*^10^. However, there were no analyses of differences in metabolic potential of microorganisms colonizing infants of different gestational age, due to the low resolution of the 16S method. A metaproteomics study revealed that gut bacteria of extremely preterm infants (< 28 weeks gestational age) produced more translation and membrane transport proteins, while the microbiomes of infants with a gestational age of 30 weeks produced more energy metabolism proteins^11^.

Genome-resolved metagenomics involves sequencing all the DNA extracted from a sample and then reconstructing genomes for the relatively abundant microorganisms present. Previous genome-resolved metagenomics studies found that the infant gut microbiome is influenced by factors such as formula feeding, the hospital room environment, and antibiotic exposure^4,12,13^. Here, we utilize genome-resolved metagenomics to analyze the effects of gestational age on the composition and metabolic potential of the premature infant gut microbiome. It is well-established that prematurity is associated with increased disease and infant mortality^14^, and this is partially due to factors involving the microbiome^15^. Understanding the effect that gestational age, i.e. the extent of prematurity, has on the microbiome may improve understanding of disease in premature infants. We found that certain bacteria occurring in infants of different gestational ages carry distinct sets of metabolic genes, thus utilizing genome-centric metagenomics to resolve how the gut microbiome is influenced by extent of prematurity.

## Results

We analyzed 900 previously reported samples from 106 premature infants with gestational ages of 24 to 32 weeks at birth **(Table S1)**. Samples were collected over the first two to three months of life. Among these infants, just 10% were classified as moderate preterm (defined as 32 to < 34-week gestation), 60% of the infants were very preterm (28 to < 32-week gestation), and 30% were extremely preterm (< 28-week gestation) (**Fig 1a)**. A correlation analysis revealed that gestational age is closely associated with birth weight (*r* = 0.84) **(Fig 1b)**, but gestational age does not display a correlation with any of the other variables in the infant metadata **(Fig 1c)**. Reconstructing genome bins from the 900 samples sequenced resulted in a dereplicated set of 1,483 genomes with an average completeness of 92% as evaluated based on the presence of bacterial single copy genes. There was no significant difference in genome completeness or sequencing depth of infants of different gestational age (**Table S2** and **Table S3**).

**Figure 1.**
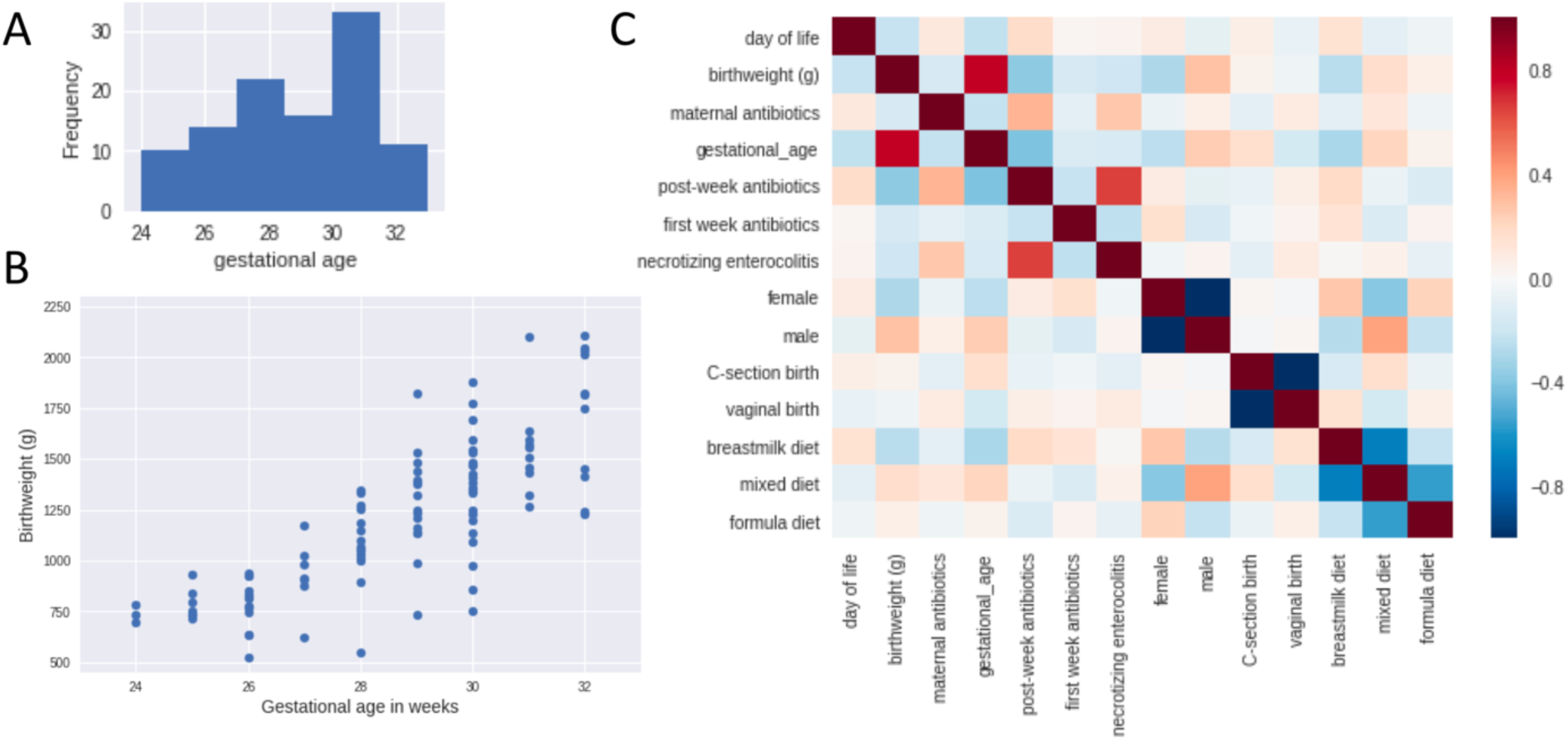
Gestational age and related infant metadata. (A) The distribution of gestational age in weeks. (B) Gestational age is correlated with birthweight (*r* = 0.84, *p*< 1 x 10^-26^, Pearson correlation). (C) Gestational age is not strongly correlated with any variable other than birthweight.

A linear regression model was applied to evaluate the effect of the infant’s characteristics as well as environmental factors on the species richness of the gut microbial community. As expected, administration of antibiotics due to a disease diagnosis after the first week of life caused a significant decrease in richness (*p* < 1 × 10^-6^, multiple linear regression analysis) **(Table 1).** The model also revealed that gestational age has a significant effect on species richness (*p* < 1 × 10^-6^, multiple linear regression analysis) **(Table 1).** To understand how gestational age’s effect on species richness changes over the course of the first few months of life, the richness of microbiomes of infants with gestational age < 28 weeks (extremely premature) was compared to that of infants with gestational age ≥ 28 weeks, at each week of life. In the first few weeks of life, microbiomes of extremely premature infants have significantly lower richness levels **(Fig 2)**. This effect is no longer present at the fifth week of life and onward.

**Table 1:** Linear regression using infant metadata variables to predict species richness.

|  | Coefficient Estimate | p-value |
| --- | --- | --- |
| intercept | -4.819817 | 0.999999 |
| infant age (days) | 0.096755 | 0.558479 |
| gestational age (weeks) | 0.385138 | 0.000000 |
| birthweight (g) | -0.001211 | 0.979418 |
| maternal antibiotics | -0.321744 | 0.215535 |
| antibiotic exposure during first week | 0.005125 | 0.915728 |
| post-week antibiotic exposure | -1.427714 | 0.000000 |
| male gender | 0.098894 | 1 |
| vaginal birth | 0.141521 | 1 |
| breastmilk | 0.566750 | 1 |
| formula | -1.014005 | 1 |
| combination of breastmilk and formula | 0.447255 | 1 |

**Figure 2.**
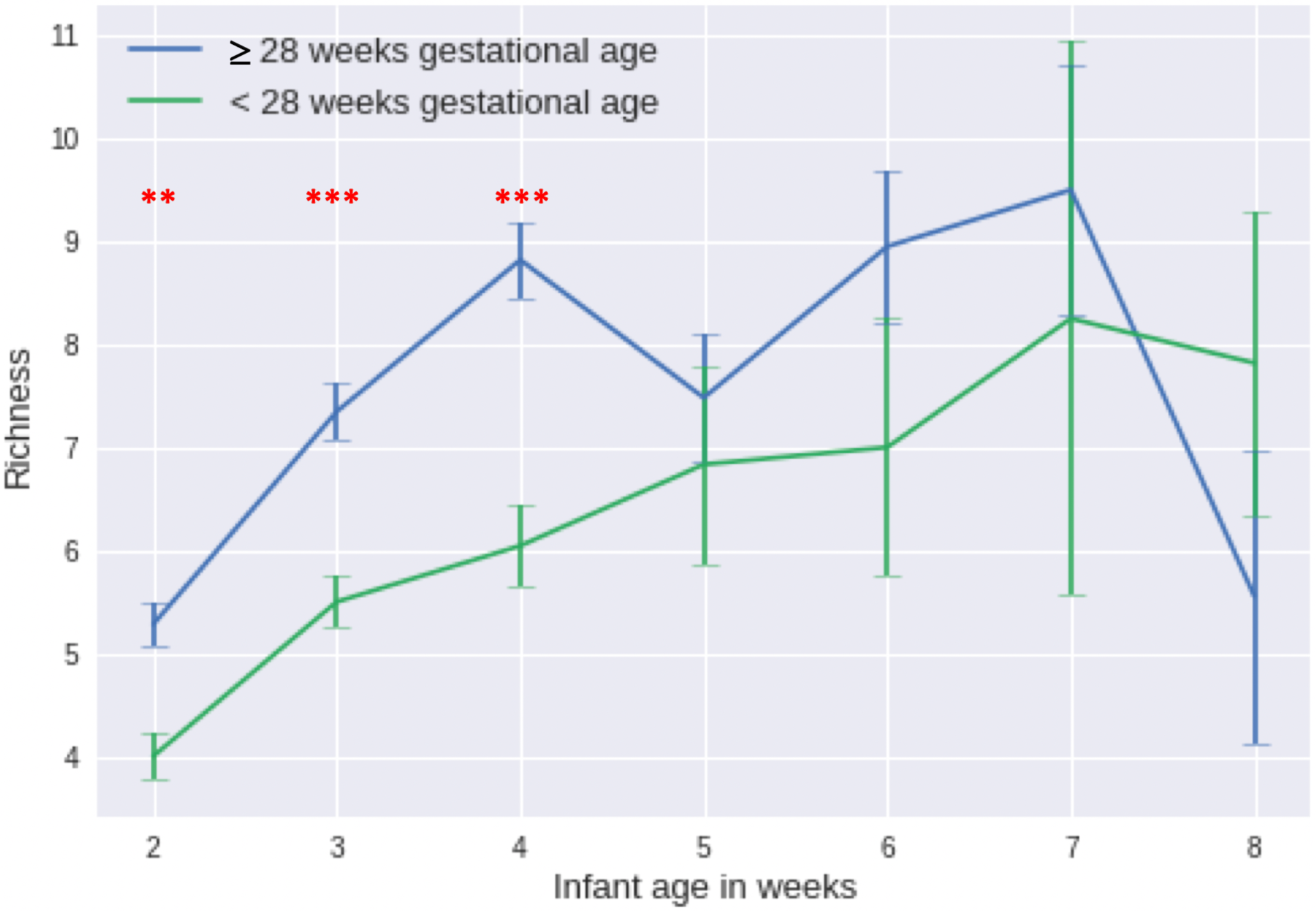
Mann-Whitney U tests were performed to compare species richness in infants with < 28 week gestational age and infants >= 28 week gestational age, at each week of life. Two asterisks indicate statistical significance at *p* < 0.005, while 3 asterisks indicate statistical significance at *p* < 0.0005. Error bars represent standard error of the mean. The number of samples in each week for infants < 28 weeks are as follows: week 2 *n* = 79, week 3 *n* = 91, week 4 *n* = 79, week 5 *n* = 13, week 6 *n* = 14, week 7 *n* = 5, week 8 *n* = 12. The number of samples in each week for infants ≥ 28 weeks are as follows: week 2 *n*= 190, week 3 *n* = 170, week 4 *n* = 85, week 5 *n* = 29, week 6 *n* = 22, week 7 *n* = 10, week 8 *n* = 10.

We compared the average taxonomic composition of microbiomes of infants with gestational age < 28 weeks and infants with gestational age ≥ 28 weeks **(Fig 3).** The figure shows small fluctuations in composition over time, but the microbiomes of individual infants can display more drastic shifts in composition. In infants with gestational age < 28 weeks, *Klebsiella* was consistently the most abundant taxa, except for during the seventh week of life where *Escherichia* was most abundant (**Fig 3a**). In infants with gestational age ≥ 28 weeks, *Escherichia* and *Klebsiella* were initially abundant but the relative abundance of these taxa appeared to decline over time (**Fig 3b**); however, this decline was not statistically significant, as the interindividual variation was substantial. When making direct comparisons of relative abundance between the two infant cohorts in the same week of life, the infants with gestational age ≥ 28 weeks had significantly higher populations of *Veillonella*, *Clostridioides* and *Clostridium* throughout the first month of life (*p* < 0.001, Mann-Whitney U test with FDR correction). When making comparisons based on corrected age, which is calculated from the time of conception to adjust for prematurity, we found no significant difference between the relative abundance values of *Veillonella* and *Clostridioides* in the two infant cohorts. The lack of difference when matching samples based on corrected age supports the hypothesis that the previously mentioned finding is due to prematurity—as the extremely premature infants reach the second month of life, they become more developmentally similar to the less premature infants, and the (likely prematurity-induced) effects that were present in early life are no longer detectable.

**Figure 3.**
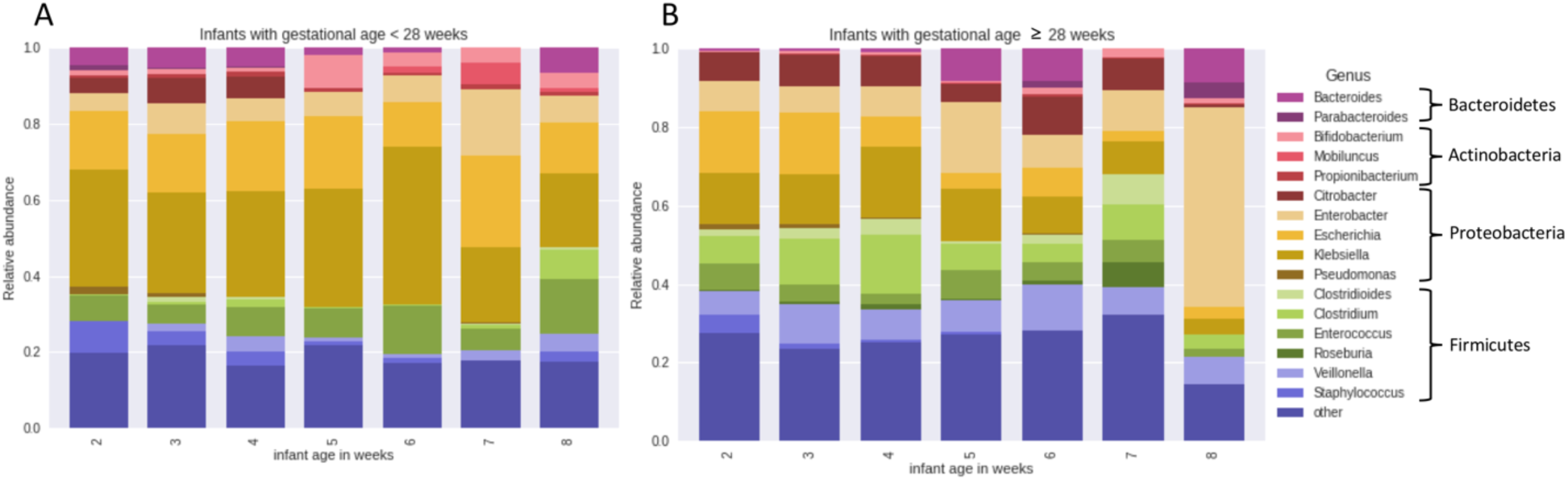
(A) The genus-level taxonomic composition of the gut community for the infants with < 28 week gestational age. The number of samples in each week for infants < 28 weeks are as follows: week 2 *n* = 79, week 3 *n* = 91, week 4 *n* = 79, week 5 *n* = 13, week 6 *n* = 14, week 7 *n* = 5, week 8 *n* = 12. (B) The genus-level taxonomic composition of the gut community for the infants with >= 28 week gestational age. The number of samples in each week for infants ≥ 28 weeks are as follows: week 2 *n* = 190, week 3 *n* = 170, week 4 *n* = 85, week 5 *n* = 29, week 6 *n* = 22, week 7 *n* = 10, week 8 *n* = 10.

We investigated the differences in metabolic potential of bacterial strains colonizing infants of varying gestational ages. Separate analyses were performed for each week of life and for each species or species group, in the case of *Clostridium*. Within each week we considered only one sample per infant to reduce bias due to resampling of the same strain in subsequent samples, allowing us to test for patterns that were consistent across infants. We uncovered trends related to the extent of prematurity, in which particular organisms exclusively colonized infants of a certain gestational age. In the second week of life, species of a novel group in *Clostridium,* all of which have genes for vitamin B biosynthesis, were only present in infants of gestational age greater than 30 weeks **(Fig 4a).** *Streptococcus salivarius*-related strains containing genes for L-Cystine transport only occurs in infants of gestational age ≤ 30 weeks **(Fig 4b).** In the third and fourth week of life, very-closely related *Enterococcus faecalis* strains with genes for the RaxABRaxC type I secretion system are present exclusively in infants of gestational age ≥ 28 weeks **(Fig 4c)**. Most of the *E. faecalis* carrying genes for this secretion system also harbor biosynthetic gene clusters for bacteriocin and lantibiotic, while none of the *E. faecalis* lacking the secretory genes were found to harbor these biosynthetic gene clusters.

**Figure 4.**
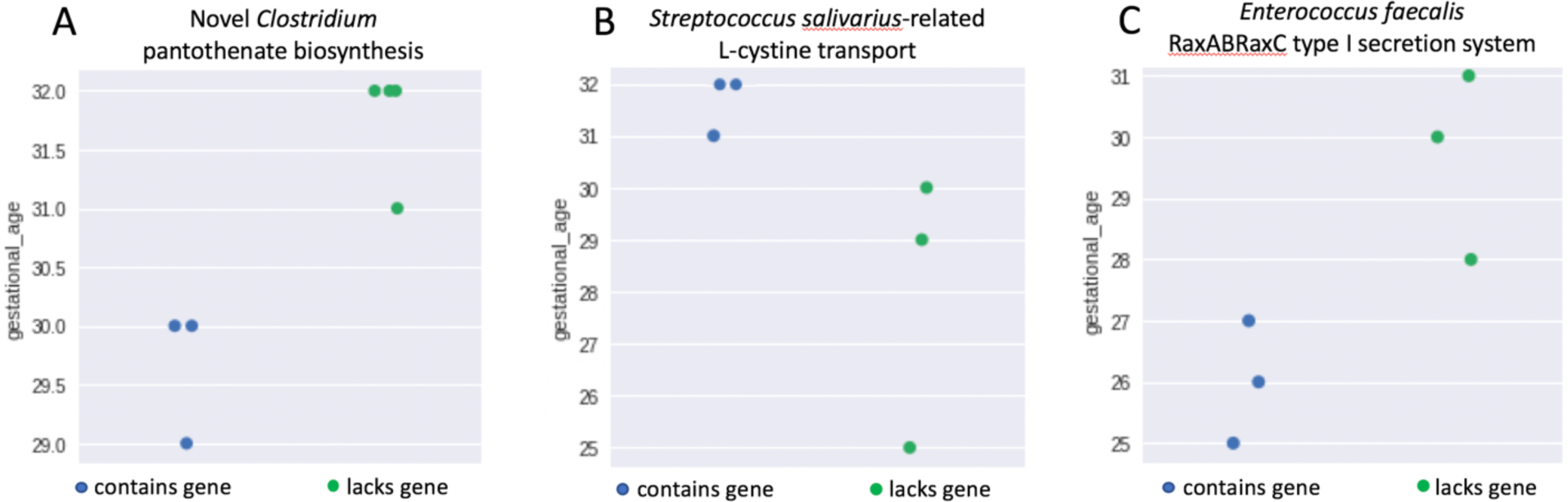
Trends in strain functional potential related to extent of prematurity. Each dot represents one genome, and the location of the dot on the y-axis indicates the gestational age of the infant in which the organism was found. Green dots indicate that the genome has the particular metabolic function listed in the title of the plot, and blue dots indicate that the genome lacks this function. (A) Genomes part of a novel group in *Clostridium* labeled as having or lacking genes for pantothenate biosynthesis. (B) *Streptococcus salivarius-*related strains labeled as having or lacking genes for an L-cystine transport system. (B) Strains of *Enterococcus faecalis* labeled as having or lacking genes for RaxAB-RaxC type 1 secretion system.

## Discussion

Our study involved analysis of the microbiomes of 107 premature infants for which a variety of metadata was collected, including each infant’s gestational age, birthweight, disease incidence, antibiotic exposure, feeding method, gender, and birth mode **(Table S1)**. Agreeing with population-based references built from historical data^16^, the gestational age of infants in this study was closely correlated with birthweight **(Fig 1b).** However, the other collected metadata did not display a correlation with gestational age **(Fig 1c)**, indicating that the findings of this microbiome study can be attributed to either (1) the gestational age, directly, or (2) differing clinical treatment among babies of varying gestational age that was not recorded in the collected metadata. This differing clinical treatment could be a particular feeding regimen (e.g., more specific than whether the infant received breastmilk, formula, or a combination), usage of a mechanical ventilator, length and timing of attachment to intravenous nutrition lines, or another factor.

Regardless of whether it is a direct or indirect effect, gestational age was strongly associated with species richness **(Table 1)**, and this effect is only present during the first month of life **(Fig 2)**. This indicates that while gestational age influences the microbes present in the few weeks immediately following birth, it does not have a persistent impact on microbiome complexity over the study period. We found that no other factors besides gestational age and post-week infant antibiotic exposure had a significant impact on richness of the gut microbiome **(Table 1).** This contrasts with findings of previous studies that associated formula feeding with increased diversity^17^ and intrapartum maternal antibiotic use with decreased diversity^18^. It is important to note, however, that in the current study, organisms can be grouped at the strain or species level, whereas prior studies mostly relied on 16S rRNA gene fragment profiling, which typically has genus-level resolution.

Perhaps the most surprising result was that the day of life (infant’s age in days) did not have a significant influence on species richness **(Table 1)**. Previous studies have shown that the microbiome of full term infants gains species over time and displays a clear increase in diversity^19^. The difference between the patterns reported here and prior studies may be largely accounted for by prematurity and, in some cases the administration of antibiotics that cause a sharp decline in microbiome diversity^20^. Because these infants were in the neonatal intensive care unit throughout our study, the consortia available to colonize them likely had lower diversity than would have been encountered in the home environment and includes bacteria considered to be hospital-associated pathogens^21^.

We found that certain organisms only colonized infants of a particular gestational age range. One such case is *Clostridium*, which are anaerobes that have been previously found in the infant gut^22^. We identified a group of *Clostridium* that was not closely related to previously sequenced organisms, and found that some organisms in this group harbor genes for biosynthesis of pantothenate, also called vitamin B₅, a water-soluble vitamin and an essential nutrient typically supplied by intestinal bacteria^23^. These novel *Clostridium* species with genes for pantothenate production were only found in less premature infants **(Fig 4a)**. It should be noted that two of the infants harboring *Clostridium* with the pantothenate biosynthesis genes are twins, born at a gestational age of 32 weeks. Genetic relatedness, exposure to the same mother’s microbiota, or other factors may have contributed to colonization by similar bacteria. However, the other infants showing gestational age-dependent colonization were unrelated. The observation that very premature infants may have comparatively lower access to vitamin B₅ than less premature infants due to strain colonization may be important because lack of pantothenic acid can adversely affect the immune system, producing a pro-inflammatory state^24^. It has been shown previously that the production of pantothenate in the gut is negatively impacted by low availability of vitamin D, which is often the case with very premature infants^25^. Thus, selection against *Clostridium* strains with the capacity for pantothenate production may be explained by increased prematurity.

*Streptococcus salivarius*-related bacteria with genes for transport of L-cystine, an amino acid essential for infants, are present only in infants of less than 31 weeks gestational age **(Fig 4b)**. The infants in this study received cysteine, which forms the cystine dimer, as part of an amino acid mixture included in the parenteral nutrition. A study evaluating plasma amino acid concentrations in infants given parenteral nutrition found that infants of lower birthweight have less of an ability to use cystine/cysteine compared to infants of higher birthweight^26^. Since the more premature infants have lower birthweight **(Fig 1b)**, the inability of the human cells to uptake cystine may lead to higher concentrations of cystine in the gut, indicating why cystine-transporting bacteria are selected for in these infants.

The trends described with *Clostridium* and *S. salivarius* both occur in the second week of life. However, *Enterococcus faecalis* displays interesting pattern in the third and fourth week of life: strains with a RaxABRaxC type I secretion system occur only in infants of greater gestational age, while strains lacking this secretion system occur exclusively in extremely premature infants **(Fig 4c).** The RaxABRaxC type I system is involved in the secretion of double-glycine-type leader peptides^27^, which occur in bacteriocins and lantibiotics^28^. All the *E. faecalis* with the RaxABRaxC type I secretion system have gene clusters for production of lantipeptides, bacteriocin, or both. In contrast, most of the *E. faecalis* lacking the secretion system (i.e., the *E. faecalis* occurring in the infants of lowest gestational age) did not have these biosynthetic gene clusters. Since bacteriocins are toxins that inhibit the growth of closely related strains, the gestational age of an infant could indirectly influence the contribution of *E. faecalis* to bacteriocin production and thus influence microbiome composition. As *E. faecalis* is a common and often abundant member of the gut microbiomes of premature infants^29^, it is possible that bacteriocin production is less common in infants of very low gestational ages.

The findings discussed above offer a strain-level perspective to what is known about how the taxonomic and functional characteristics of the gut microbial community change depending on the infant’s gestational age^10,11^. By analyzing the metabolic potential of each genome, we found evidence that the extent of prematurity, either directly or indirectly, can affect the gut microbiome. Given evidence that a lower gestational age may limit bacteriocin and vitamin production, which are factors that can impact community structure and lead to inflammation^30,31^, these findings may inform our understanding of diseases associated with dysbiosis of the microbiome, especially in very premature infants.

## Methods

Sample collection and metagenomic data processing for these samples were previously described^32^. Briefly, fecal samples were collected from premature infants residing in the neonatal intensive care units (NICU) of the Magee-Women’s Hospital in Pittsburgh, PA, and a PowerSoil DNA isolation kit (Mo Bio Laboratories, Carlsbad, CA) was used to extract the DNA, which was then sequenced on an Illumina platform (further details available here^21,33,34^). The samples analyzed in this study have been previously reported, and the reads are publicly available at NCBI as described here^12,21,33,34^.

The reads were trimmed using Sickle (https://github.com/najoshi/sickle) and cleared of human contamination through read mapping with Bowtie2^35^. IDBA-UD^36^ was used to assemble the reads of each sample and was also used to generate co-assemblies by combining the reads of all the samples from a particular infant. The genes on the scaffolds were predicted using Prodigal^37^. The scaffolds were grouped into genome bins using concoct^38^ and redundant bins were dereplicated using dRep^39^ v0.4.0. Centrifuge^40^ was used for initial assignment of taxonomy, which was then verified by confirming the phylogenetic profiles of the scaffolds in the genome bin. When majority of the scaffolds in a genome bin were of unknown phylogenetic profile, the organism was concluded to be novel. For annotation of metabolic capabilities, the sequences were searched against the Kyoto Encyclopedia of Genes and Genomes (KEGG)^41^ using profile hidden Markov models, and the results were used to generate a KEGG metabolism profile for each organism that displayed the fraction of each KEGG module encoded by that genome. For specialized identification of biosynthetic gene clusters, antiSMASH^42^ was used to annotate genes from particular scaffolds of interest. Organisms were considered to be present in a particular sample if the genome bin showed breadth of coverage over 99%.

When calculating correlations, Pearson’s product-moment correlation was used for two continuous variables and the point biserial correlation was used for pairs that contained at least one categorical variable. To model the effects of infant characteristics and clinical treatments on gut species richness levels, linear regression from the scikit-learn^43^ package was utilized. Mann-Whitney U tests were performed for comparison of relative abundance values and for the comparison of richness values at each week, and p-values were corrected using the False Discovery Rate (FDR) method. Spearman correlations were performed to identify trends over time. When carrying out the strain-focused portion of the study, separate analyses were performed for each week of life and for each species, to remove effects of these factors. Only one sample per infant was included in each analysis to avoid bias due to repeated samples. When a genome was considered to be lacking a particular gene, the entire dRep set (all genomes belonging to the same secondary cluster in dRep) was checked to ensure that another genome in that set did not harbor the gene of interest. Associations were considered spurious and removed if the validation step did not confirm the findings.

## Author Contributions

JFB and MJM conceived of the project. MRO processed the sequence data. SFR performed the data analysis and interpreted the results. SFR and JFB wrote the manuscript, and all authors contributed to manuscript revisions.

## Acknowledgements

We acknowledge Robyn Baker for recruiting infants, Brian Firek for performing DNA extractions, Christopher Brown for scripts to calculate genome coverage, and David Burstein for the KEGG HMM annotation pipeline.

